# Replating induces mTOR-dependent rescue of protein synthesis in Charcot-Marie-Tooth diseased neurons

**DOI:** 10.1101/2025.08.26.672367

**Authors:** Julianna Koenig, Alexys McGuire, Yara Homedan, Jessica Alberhasky, Daniel W. Summers

**Author notes:** Address Correspondence to: Daniel Summers, 129 E. Jefferson St. Iowa City, IA 52242, Contact.

## Abstract

Charcot-Marie-Tooth disease (CMT) is an inherited peripheral neuropathy characterized by sensory dysfunction and muscle weakness, manifesting in the most distal limbs first and progressing more proximal. Over a hundred genes are currently linked to CMT with enrichment for activities in myelination, axon transport, and protein synthesis. Mutations in tRNA synthetases cause dominantly inherited forms of CMT and animal models with CMT-linked mutations in these enzymes display defects in neuronal protein synthesis. Rescuing protein synthesis in CMT mutant neurons could offer exciting therapeutic options beyond symptom management. To address this need, we expressed CMT-linked variants in tyrosyl tRNA synthetase (YARS-CMT) in primary sensory neurons and evaluated impacts on protein synthesis and cell viability. YARS-CMT expression reduced protein synthesis in these neurons prior to the onset of caspase-dependent axon degeneration and cell death. To determine how YARS-CMT expression affects axon outgrowth, we dissociated and replated these neurons to stimulate axon regeneration. To our surprise, axonal regrowth occurred normally in replated YARS-CMT neurons. Moreover, replating YARS-CMT neurons rescued protein synthesis. Inhibiting mTOR suppressed rescue of protein synthesis after replating, consistent with its significant role in protein synthesis during axon regeneration. These discoveries identify new avenues for augmenting protein synthesis in diseased neurons and restoring protein synthesis in CMT or other neurological disorders.

## INTRODUCTION

Neurons of the peripheral nervous system (PNS) are responsible for relaying somatosensory information to the central nervous system (CNS) and stimulating muscle fibers. Both PNS and CNS neurons are post-mitotic and must sustain these vital functions for an organism’s entire lifespan. Neurons establish connections throughout the body via long axons, extending more than a meter in the human PNS. This extreme length coupled with a high metabolic demand renders axons especially vulnerable to damage and stress (Coleman and Hoke, 2020). As such, peripheral neuropathies are characterized by progressive and sometimes irreversible damage to the PNS, causing severe disability and diminished quality of life (Hanewinckel et al., 2016). With treatment options focused on symptom management, understanding the mechanistic basis of peripheral neuropathies will identify new opportunities for halting disease progression or even reversing PNS damage.

Charcot-Marie-Tooth disease (CMT) is the most common inherited peripheral neuropathy with an estimated prevalence of 1 in 2,500 to 1 in 10,000 predicted from epidemiological studies (Skre, 1974; Martyn and Hughes, 1997; Barreto et al., 2016). CMT is characterized by progressive distal muscle weakness and sensory loss arising from the gradual dysfunction of peripheral axons through demyelination or intrinsic degeneration. These pathological mechanisms form the basis of CMT categorization into demyelinating (CMT1) or axonal (CMT2) forms, with disorders exhibiting features of both classified as intermediate CMT. Further subcategorization within each form is defined by the causative mutation(s) (Pareyson and Marchesi, 2009).

Aminoacyl-tRNA synthetases (aaRS) represent the largest gene family implicated in CMT, contributing to both CMT2 and dominant forms of intermediate CMT (DI-CMT) (Zhang et al., 2021). AaRS covalently link tRNAs with their cognate amino acid, thereby charging tRNAs for polypeptide synthesis at the ribosome. CMT-linked mutations predominantly occur in the aaRS catalytic domain suggesting loss of aminoacylation activity as the underlying cause, however there are notable exceptions and discrepancies reported for most aaRS (Zhang et al., 2021). For example, autosomal dominant mutations in tyrosyl-tRNA synthetase (YARS) cause DI-CMTC yet there is a disconnect between aminoacylation activity and neuropathy. Of the three most studied CMT-linked variants in YARS, G41R and 153-156delVKQV decrease aminoacylation while E196K displays normal aminoacylation, however all three provoke neuropathy in Drosophila and mouse models (Jordanova et al., 2006; Storkebaum et al., 2009; Froelich and First, 2011). Gain-of-function interactions for these CMT variants may also contribute to neuropathy beyond directly impacting protein synthesis (Blocquel et al., 2017; Bervoets et al., 2019; Ermanoska et al., 2023; Rhymes et al., 2024).

Despite variable effects on aminoacylation activity, Drosophila models of CMT- YARS report reductions in global translation from motor and sensory neurons expressing any of the above-mentioned YARS variants (Niehues et al., 2015). Moreover, the integrated stress response (ISR) is hyperactivated in CMT-YARS mouse models and ISR inhibition alleviates neuropathy phenotypes. (Spaulding et al., 2021; Zuko et al., 2021). These studies collectively point to significant roles for protein homeostasis in aaRS-CMT neuropathies, yet there remain important gaps in knowledge. For example, protein synthesis is critical for axon outgrowth during neurodevelopment, yet symptom onset in patients with aaRS-CMT variants usually occurs during late childhood or early adolescence when PNS axons have already innervated their targets.

In this study we identify protein synthesis defects induced by CMT mutations in tyrosyl-tRNA synthetase (YARS) using a primary sensory neuron model. Protein synthesis defects preceded the onset of axon degeneration and caspase activation. We used a replating procedure that induces axon regeneration to assess how these CMT- YARS variants affect axon regrowth. While chemical inhibition of protein synthesis suppressed axonal regrowth, we were surprised to observe normal axon regrowth in CMT-YARS neurons. Moreover, replating reversed protein synthesis defects in CMT- YARS neurons through mTOR signaling and new transcription. We propose that developmental outgrowth pathways triggered during replating protect neurons from CMT-YARS and that these safeguards gradually diminish as PNS neurons mature, leading to disease onset.

## RESULTS

### CMT mutations in tyrosyl-tRNA synthetase (YARS) provoke sensory neuron death and axon degeneration

Autosomal dominant mutations in human YARS cause DI-CMTC and introducing YARS-CMT variants into a wildtype background is sufficient to induce peripheral neuropathy in most model systems (Niehues et al., 2015; Bervoets et al., 2019; Spaulding et al., 2021; Zuko et al., 2021; Hines et al., 2022; Rhymes et al., 2024). For this study we used embryonic-derived mouse sensory neurons isolated from dorsal root ganglia (DRGs). These neurons are readily manipulated with lentiviruses and regrow severed axons which enabled us to assess how YARS-CMT variants impact axon regrowth. We prepared lentivirus expressing human YARS constructs downstream of the ubiquitin promoter. An empty vector and wildtype YARS were used as controls. We generated lentiviruses expressing three different variants identified in DI-CMTC populations, G41R, D81I, and 153-156delVKQV(del153) (Jordanova et al., 2003; Jordanova et al., 2006; Hyun et al., 2014). We intended to include another well-studied variant, E196K, however we consistently observed poor protein expression and did not pursue this variant further.

We first evaluated whether prolonged expression of CMT-YARS variants affected axon integrity. Lentiviral preparations were applied to cultured DRGs at 2 days *in vitro* (DIV). On DIV 7, the degree of axon degeneration was quantified using an ImageJ macro that measures particle circularity to delineate fragmented versus intact axon area to calculate a “degeneration index” from 0 to 1 (least to most degeneration, respectively) (Gerdts et al., 2011). DRG neurons expressing all YARS variants displayed significant axon degeneration compared to empty vector control (Fig. 1*A*). Importantly, expression of WT-YARS did not differ from the vector condition, indicating that elevated levels of tyrosyl-tRNA synthetase did not underlie the observed axon degeneration (Fig. 1*A*). At the same timepoint, we also observed widespread cell death (greater than 60%) in all three mutant conditions, as measured by incorporation of the membrane impermeable dye, NucSpot 470 (Fig. 1*B*).

**Figure 1.**
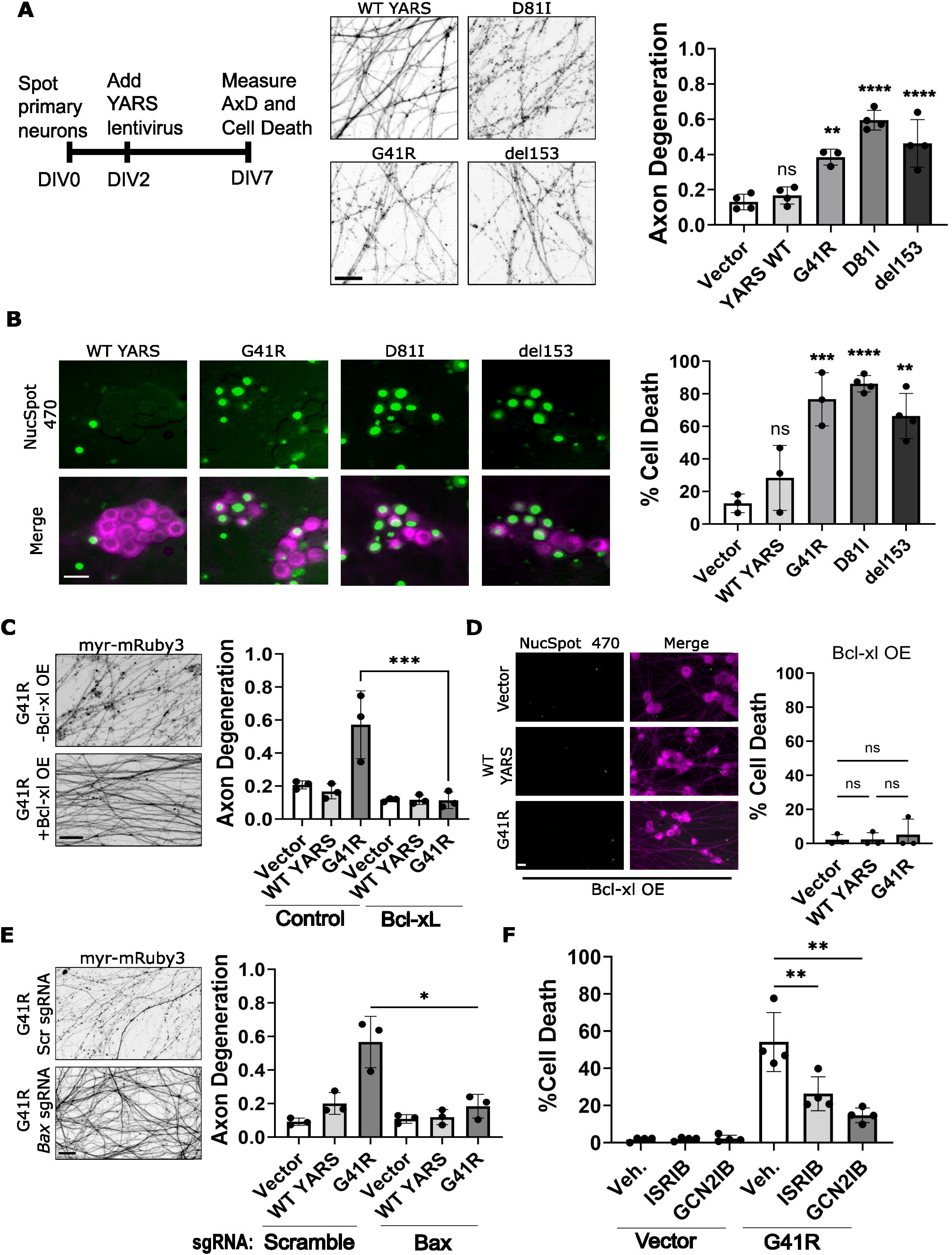
CMT-linked variants in tyrosyl-tRNA synthetase (YARS) induce caspase-dependent cell death and axon degeneration. **(A)** Primary sensory neurons from embryonic dorsal root ganglia (DRG) were transduced with lentivirus expressing the indicated YARS variant on DIV 2. Axon fragmentation was apparent by DIV 7 from neurons expressing YARS variants with CMT-linked mutations. Example images and quantification from 3-4 independent cultures are shown on the right. **(B)** YARS-CMT expressing neurons also stain positive for the cell-impermeable dye NucSpot which enters cells with compromised membrane permeability and fluoresces in complex with DNA in the nucleus. Example images and quantification (N = 3-4) are shown on the right. Cells were transduced with YARS-expressing lentivirus and a lentivirus expressing the anti-apoptotic protein Bcl-xL. Expressing Bcl-xL suppressed axon degeneration **(C)** and cell death **(D)** (N = 3). Crispr-inactivating the proapoptotic protein BAX suppressed axon degeneration **(E)** in G41R-expressing neurons compared to scrambled (scr) sgRNAs (N = 3). **(F)** Applying ISRIB (500nM) or GCN2IB (10µM) suppressed cell death in G41R-YARS expressing neurons (N = 3). Scale bar = 20µm. Error bars represent +/-1 SD. For statistical tests, one-way ANOVA was performed with post-hoc t-tests where *p<0.05, ** p<0.01, ***p<0.005, and **** p<0.001. ns. not significant.

Caspase-dependent apoptosis promotes neuronal loss during development and in the context of neurodegeneration. Inhibiting caspase activation by Bcl-xL overexpression or CRISPR inactivating BAX suppressed cell death and axon degeneration in neurons expressing YARS G41R (Fig. 1*C-E*) as well as D81I-YARS and del153-YARS (Supplementary Fig. 1 *A&B*). Bcl-xL expression did not affect CMT-YARS protein levels (Supplementary Fig. 1C) indicating neuroprotective effects were due to this protein’s antiapoptotic function rather than downregulation of mutant CMT-YARS constructs.

The ISR is activated in spinal motor neurons and DRGs from CMT mouse models possessing CMT mutations in glycyl-tRNA synthetase (Spaulding et al., 2021; Rhymes et al., 2024). The motor neuropathy present in these CMT mouse models is reduced in a GCN2^-/-^ background, suggesting a contribution of the ISR to CMT pathology (Spaulding et al., 2021). To evaluate whether the ISR promotes neurodegeneration in our primary neuron model, we administered two different small molecule inhibitors targeting the ISR, ISRIB or GCN2iB, one day after applying CMT- YARS lentivirus and quantified cell death. Both inhibitors suppressed cell death induced by G41R-YARS (Fig. 1*F*) confirming a prodegenerative role of the ISR in this CMT model.

### CMT-YARS reduces protein synthesis in DRG sensory neurons

We next evaluated whether CMT-YARS variants affect protein synthesis in sensory neurons. We overexpressed Bcl-xL via lentiviral transduction to prevent caspase activation and circumvent changes in protein synthesis expected downstream of apoptosis. YARS variants G41R and del159 reduce aminoacylation activity *in vitro* and *in vivo* (Niehues et al., 2015) while the D81I variant has not been determined. We used two strategies to evaluate protein synthesis in DRG neurons expressing these variants. We first labeled newly synthesized proteins with the amino acid analog O-propargyl-puromycin (OPP) then used Click Chemistry to visualize this analog with the fluorescent probe AZdye 488. DRGs transduced with G41R or D81I YARS showed a modest decrease in OPP fluorescence compared to empty vector controls, with statistically significant differences only observed between YARS D81I and the empty vector condition (Fig. 2*A*). The translation inhibitor cycloheximide did not completely suppress OPP fluorescence, indicating off-target fluorophore labeling during the click chemistry reaction which would decrease sensitivity of the assay. Additionally, OPP labeling requires cell fixation, limiting our investigation to a single timepoint and preventing analysis of the temporal dynamics of protein synthesis.

**Figure 2.**
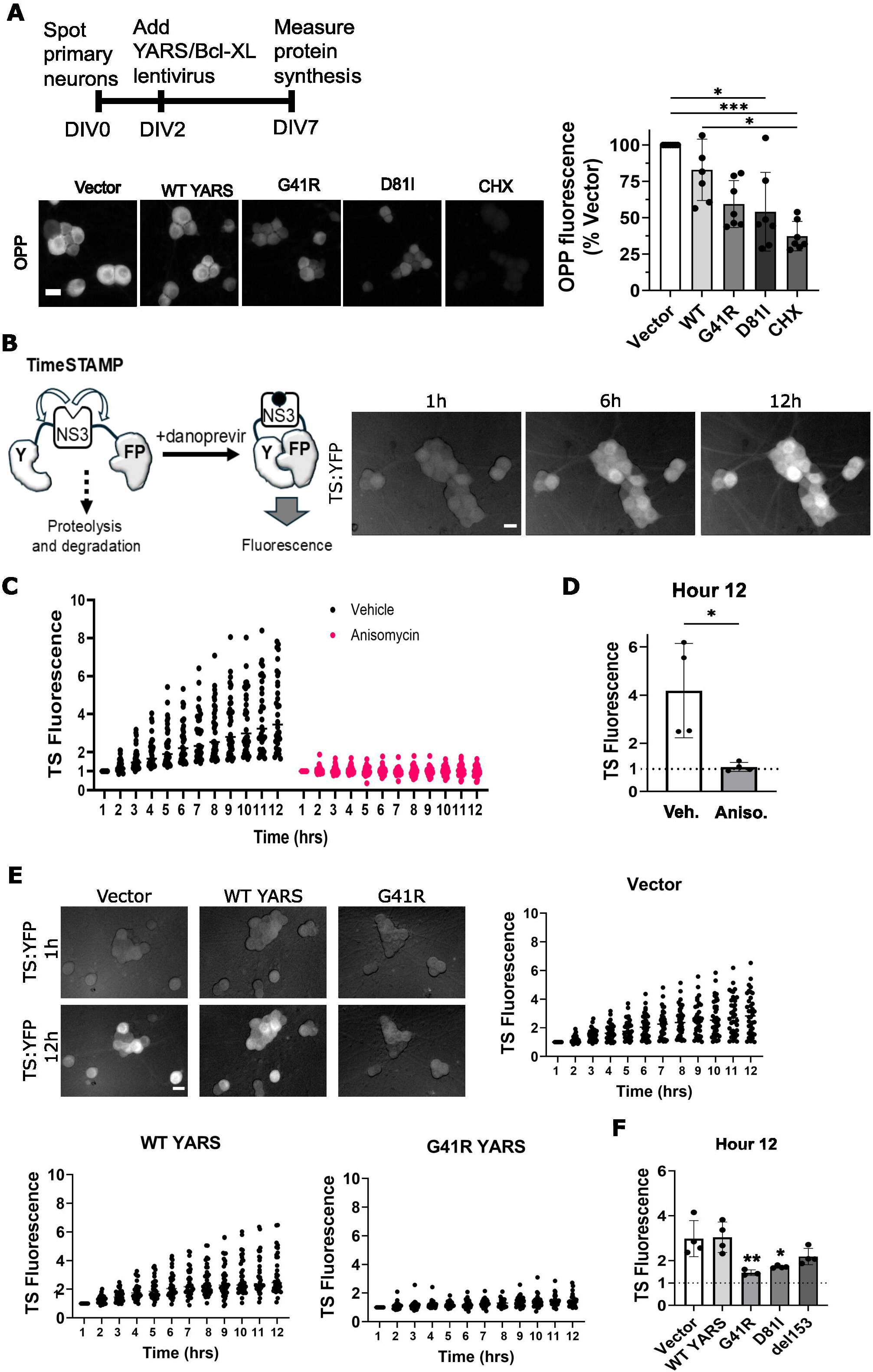
CMT-YARS expression reduces protein synthesis. DRG sensory neurons were transduced on DIV2 with the indicated YARS lentivirus and Bcl-xL lentivirus to suppress caspase activation. **(A)** Newly synthesized proteins were labeled with O-propargyl-puromycin (OPP) for 10min and visualized after Click Chemistry with AZdye488. OPP fluorescence was reduced in the presence of G41R or D81I YARS compared to empty vector or wildtype YARS. Applying the protein synthesis inhibitor cycloheximide (CHX) prior to labeling further reduced protein synthesis. Representative images and quantification (N=6) are shown. **(B)** Diagram of TimeStamp (TS) reporter for newly synthesized proteins. Steady state TS fluorescence is low as newly synthesized reporter undergoes proteolysis and protein degradation. Applying the NS3 inhibitor danoprevir stabilizes newly synthesized TS protein, split YFP fragments assemble, and fluorescence increases over time. On the right, representative images show TS fluorescence after danoprevir application. **(C)** Quantification of TS fluorescence in individual cells across 4 experimental replicates after danoprevir addition as a ratio of 1hr fluorescence. In **(D)** the change in fluorescence at twelve hours was averaged in four independent experiments (from at least forty cells). Applying the translation inhibitor anisomycin (10 µM) suppresses TS fluorescence after danoprevir treatment **(C)** & **(D)**. **(E)** G41R YARS reduces TS fluorescence after danoprevir treatment compared to an empty vector or WT YARS (N=4) with 12-hour time points shown in **(F)** from experiments performed with all three CMT-YARS variants. Scale bar = 20µm. Error bars represent +/-1 SD. A Kruskal-Wallis analysis was performed for data shown in **(A)** with Dunn’s post-hoc test for multiple comparisons where *p<0.05 and ***p<0.005. For experiments in (**D)** & **(F)**, a one-way ANOVA was performed with post-hoc t-tests where *p<0.05, ** p<0.01, and ***p<0.005.

As an alternative approach to measure protein synthesis in live cell cultures, we leveraged a genetically encoded fluorescent reporter for protein synthesis called TimeSTAMP (TS) (Lin et al., 2008; Lin and Tsien, 2010). The TS construct consists of the nonstructural protein 3 (NS3) protease flanked by two linker regions and split yellow fluorescent protein (YFP) fragments. The protein reporter is continuously synthesized and degraded, as the NS3 protease targets cleavage sites in the adjacent linker regions. Applying an NS3 protease inhibitor (danoprevir) prevents degradation of newly synthesized TS protein, assembly of the split YFP fragments, and subsequent change in fluorescence to serve as a readout for new protein synthesis (Fig. 2*B*). DRGs were transduced with lentivirus expressing TS on DIV 5, and danoprevir added on DIV 7.

Over the next 12 hours, fluorescence in individual neurons was tracked once an hour with an automated microscope, then each neuron’s fluorescence signal was normalized to its value at hour 1 post danoprevir treatment (Fig. 2*C*). Fluorescence intensity increased over time however the rate varied from neuron-to-neuron (Fig. 2*C*), consistent with other single-cell studies of protein synthesis (Han et al., 2014). We limited our statistical analysis to the relationship between conditions at the final timepoint with relative fluorescence intensities averaged from at least ten cells per experimental replicate (evaluating at least 30 total cells per condition). Importantly, treatment with the translation inhibitor anisomycin suppressed the increase in fluorescence, confirming this signal represents newly synthesized TS (Fig. 2*C* & *D*).

DRGs expressing any CMT-YARS variant displayed substantial defects in protein synthesis at DIV 8 compared to Vector and WT-YARS conditions (Fig. 2*E & F* and Supplementary Fig. S2*A & B*). Del153 YARS expression caused a milder synthesis defect, with severity falling between negative controls and G41R/D81I YARS (Supplementary *Fig*. S2*B*). The decrease in TS fluorescence was confirmed by western blot from whole cell extracts for the TS construct (Supplementary Fig. S2*C*).

We hypothesized the protein synthesis defect on DIV 8 is an early outcome of CMT-YARS expression and precedes onset of apoptosis. To evaluate this further, we measured protein synthesis on DIV 5, three days after viral transduction (Fig. 3*A*). Even at this early timepoint, G41R-YARS expression caused a protein synthesis defect compared to empty vector and WT-YARS controls (Fig. 3 *B-D*). In the absence of Bcl-xL we observe no axon degeneration at this timepoint (Fig. 3*E*) and caspase 3/7 activity was detected in 12% of G41R expressing neurons, lower than the 40% caspase3/7 positive neurons observed two days later (Fig. 3*F*). Therefore, G41R-induced defects in protein synthesis occur days before axon fragmentation and widespread induction of caspase activation.

**Figure 3.**
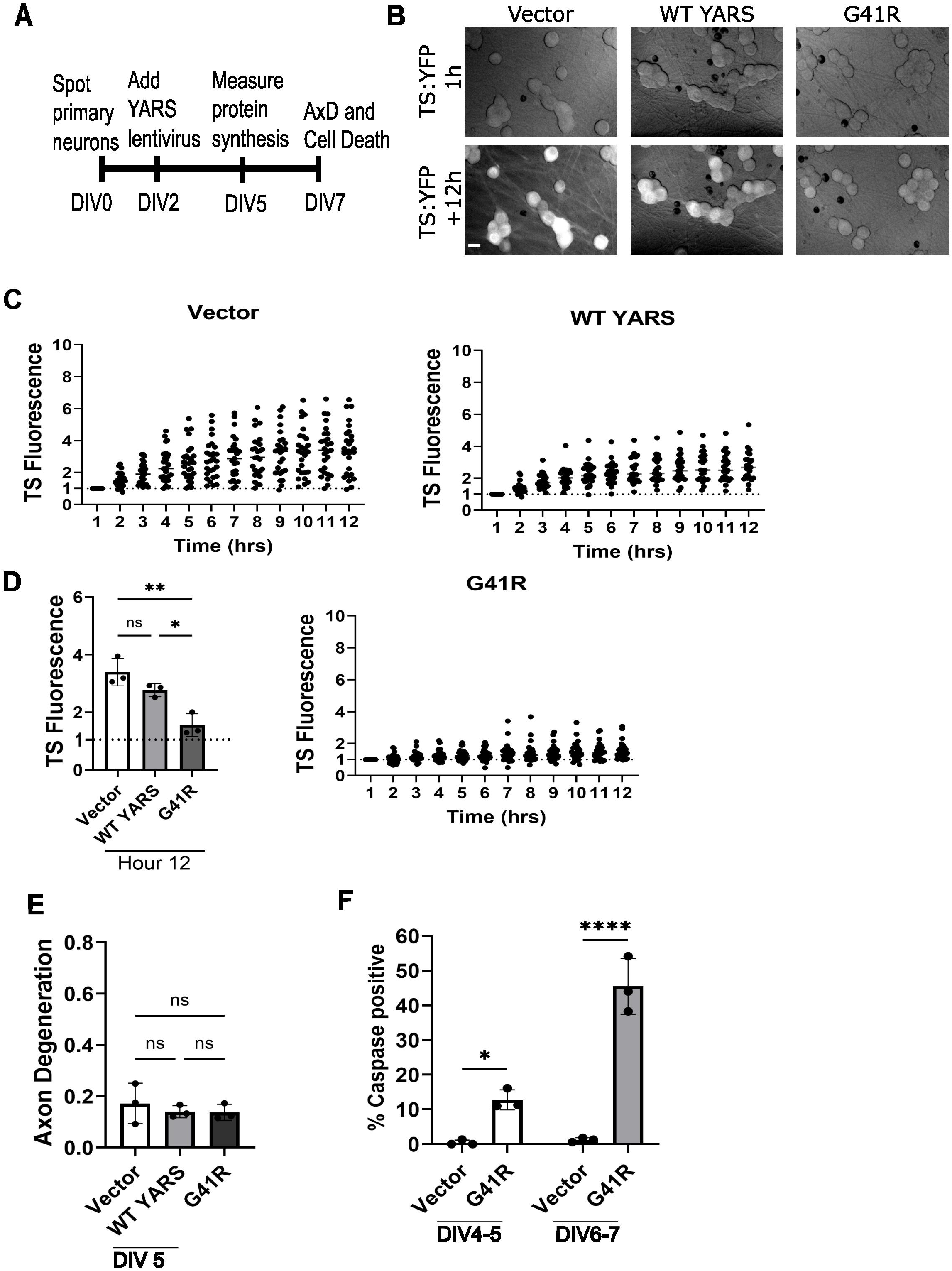
Reduced protein synthesis precedes indicators of neurodegeneration in neurons expressing G41R-YARS. **(A)** We measured protein synthesis in neurons expressing G41R-YARS on DIV5. **(B) & (C)** TS fluorescence after danoprevir addition in neurons transduced with lentivirus expressing G41R-YARS, WT YARS, or an empty vector over a 12-hour time period with TS fluorescence at 12 hours in **(D)** (N=3). **(E)** Axon degeneration on DIV 5. **(F)** We visualized caspase3/7 activation over a 12-hour time period with a cell-permeable fluorescent dye that translocates to the nucleus and fluorescence in the presence of activated caspases. We noted positive caspase labeling was transient in a dying population, so we tracked the total number of caspase-positive cells over a 12-hour interval. A small, though significant population of caspase-positive cells were detected in the presence of G41R YARS between DIV 4 and DIV 5. This percentage increased between DIV 6 and DIV7. Scale bar = 20µm. Error bars represent +/-1 SD. For statistical tests, one-way ANOVA was performed with post-hoc t-tests where *p<0.05, ** p<0.01, and ***p<0.005.

### Investigating effects of CMT-YARS on axon regeneration

Axon outgrowth during development and regeneration depends on new protein synthesis. Introducing CMT-YARS by lentivirus on DIV 2 precluded us from assessing its effects on early axon outgrowth during DIV 0-1. To circumvent this limitation, we utilized a replating protocol to remove axons and stimulate axon regeneration (Frey et al., 2015) (Fig. 4*A*). CMT-YARS and Bcl-XL were transduced into DRG sensory neurons as described above. On DIV 8, DRG cultures were briefly trypsinized to detach cells from the plate. The collected DRG suspensions were triturated and resuspended to shear off the axons, then reseeded onto a new plate. Replated DRGs sprout new axons with growth cones within the first few hours after replating (Fig. 4*A*). Consistent with prior studies activating adenylate cyclase with forskolin, applying a cell permeable cyclic-AMP analog accelerated axon regeneration (Kilmer and Carlsen, 1984; Cai et al., 1999; Neumann et al., 2002; Qiu et al., 2002; Ghosh-Roy et al., 2010; Frey et al., 2015; Hao et al., 2016) (Fig. 4*B-D*). Conversely, anisomycin treatment inhibited regeneration, confirming axon regrowth following replating is dependent on new protein synthesis (Twiss and Shooter, 1995). Similar to TS studies in Figure 2 and Figure 3, we limited our statistical analysis to the relationship between conditions at the final timepoint of 11 hours post replating (hpr) (Fig. 4*D*). Unexpectedly, the rate of axon regrowth from replated DRGs expressing G41R was similar to regrowth from empty vector and WT controls (Fig. 4*E-G*). We tracked axonal growth up to 36 and 84 hpr and observed a modest, non-significant decrease in G41R-expressing cells at 84 hpr (Fig. 4*H* & *I*).

**Figure 4.**
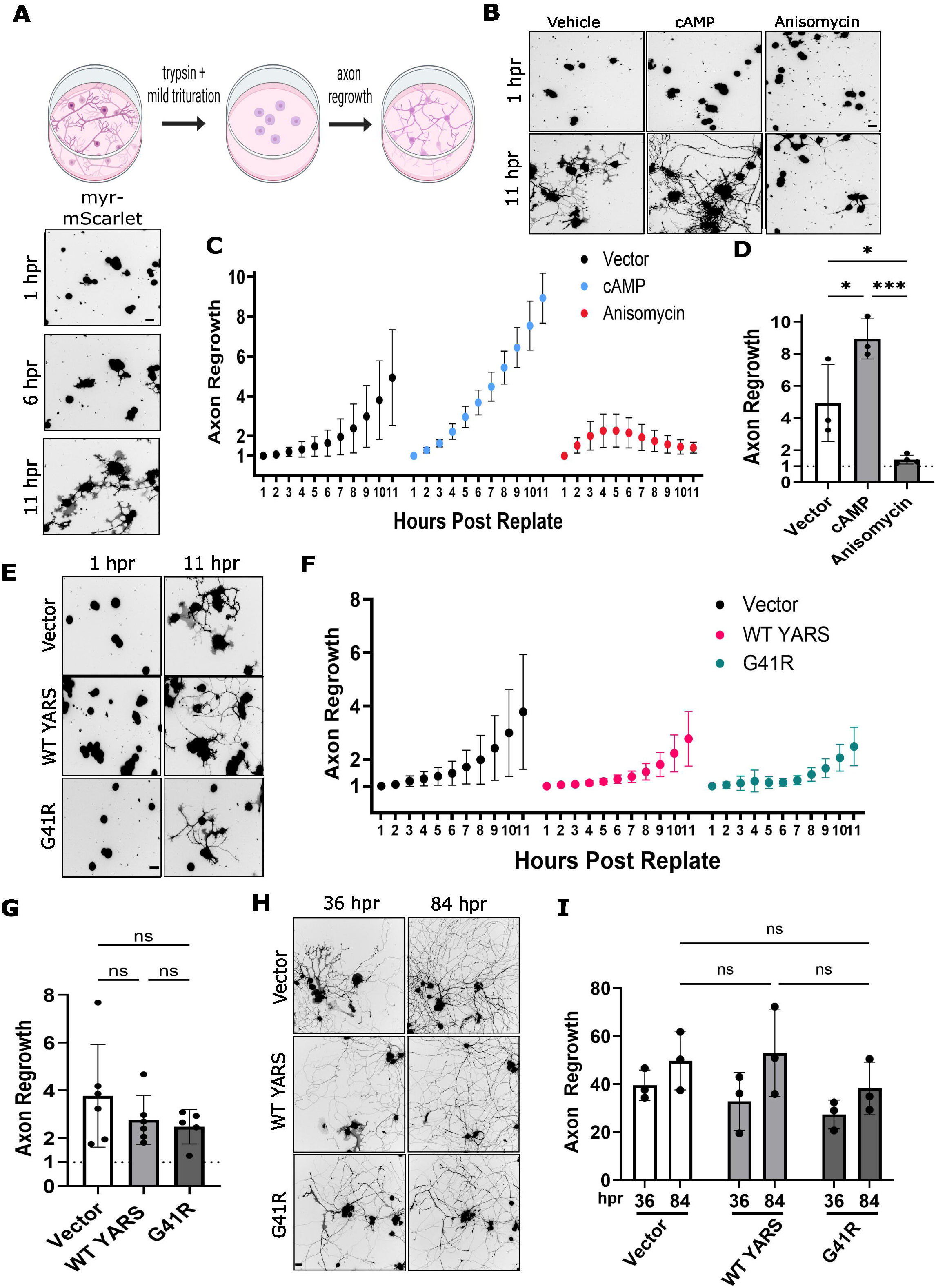
G41R-YARS does not impair axon regeneration after replating. **(A)** Sensory neurons were transduced with YARS lentiviruses as described in prior experiments as well as Bcl-xl expressing lentiviruses to suppress caspase-dependent cell death. On DIV 8, neurons were lifted from the plate with trypsin digestion, axons sheared off by mild trituration, and neurons seeded onto new plates. Axonal regrowth was visualized over eleven hours with automated microscopy and quantified by measuring total axon area at a given timepoint normalized to total axon area at time 1hr. Example images are shown below. **(B) & (C)** Applying a cell permeable cAMP analog (50 µM) accelerated axon regrowth while the protein synthesis inhibitor anisomycin (10 µM) suppressed axon regrowth. **(D)** Axon regrowth in neurons treated with cAMP or anisomycin at eleven <u>h</u>ours <u>p</u>ost replating (11hpr) (N=3). **(E) – (G)** Axon regrowth in G41R-YARS neurons was unchanged compared to WT-YARS or an empty vector with eleven hours post replating (11 hpr) (N = 5-6). **(H) & (I)** Axon regrowth in G41R-YARS neurons was similar to control conditions at 36 hpr. A slight reduction in axon area was noted at 84 hpr in G41R-YARS neurons (N = 3). Scale bar = 40µm. Error bars represent +/-1 SD. For statistical tests, one-way ANOVA was performed with post-hoc t-tests where *p<0.05, ** p<0.01, and ***p<0.005. Replating diagram was generated with BioRender.

Normal axon regeneration in G41R-expressing neurons prompted us to investigate new protein synthesis. We measured TS fluorescence immediately after replating over a twelve-hour period. To our surprise, protein synthesis in replated, G41R-expressing neurons was unchanged compared to empty vector and WT-YARS (Fig. 5*A* & *B*). We suspected rescue of protein synthesis might subside over time as replated neurons mature. We tested this prediction by incubating replated neurons for 84 hours before adding danoprevir and measuring TS fluorescence over the next 12 hours (96 hpr). While TS synthesis in the G41R condition initially appeared normal compared to vector and WT-YARS, synthesis tapered off and by 96 hpr was significantly reduced compared to empty vector and WT-YARS (Fig. 5*C* & *D*). Therefore, replating temporarily alleviates protein synthesis defects present in G41R-YARS neurons.

**Figure 5.**
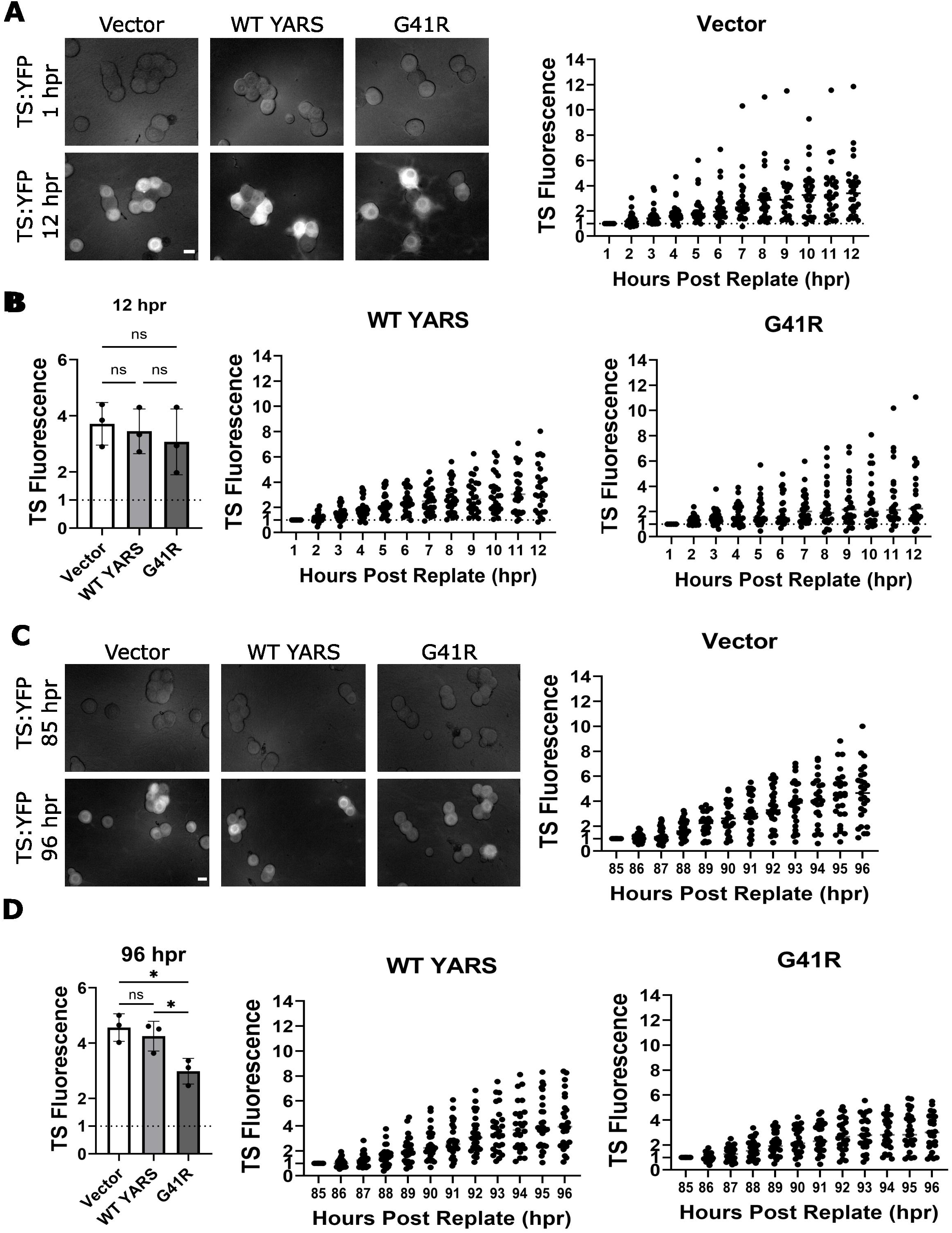
Replating rescues protein synthesis in G41R-YARS neurons. **(A)** TS fluorescence was measured in replated neurons transduced with lentivirus expressing G41R-YARS, WT-YARS, or an empty vector. **(B)** TS Fluorescence at twelve-hours post danoprevir application in replated neurons (N= 3). **(C)** Neurons eighty-four hours after replating were treated with danoprevir and TS fluorescence tracked over twelve hours with terminal fluorescence at ninety-six hours post-replating shown in **(D)** (N = 3). Scale bar = 20µm. Error bars represent +/-1 SD. For statistical tests, one-way ANOVA was performed with post-hoc t-tests where *p<0.05, ** p<0.01, and ***p<0.005.

We examined protein levels of endogenous YARS and lentiviral-expressed G41R-YARS after replating to ascertain whether changes in TS synthesis were due to alterations in protein levels. For example, replating might upregulate endogenous YARS and suppress deleterious effects of G41R-YARS. We first compared endogenous YARS in pre-replated samples to those collected immediately after replating (Figure 6*A*) or samples collected 24hpr (Figure 6*B*). Endogenous YARS protein did not change between either timepoint. Endogenous YARS appeared to decrease 24hpr, however total protein levels were also reduced in these samples possibly due to cell death occurring during the replating procedure.

**Figure 6.**
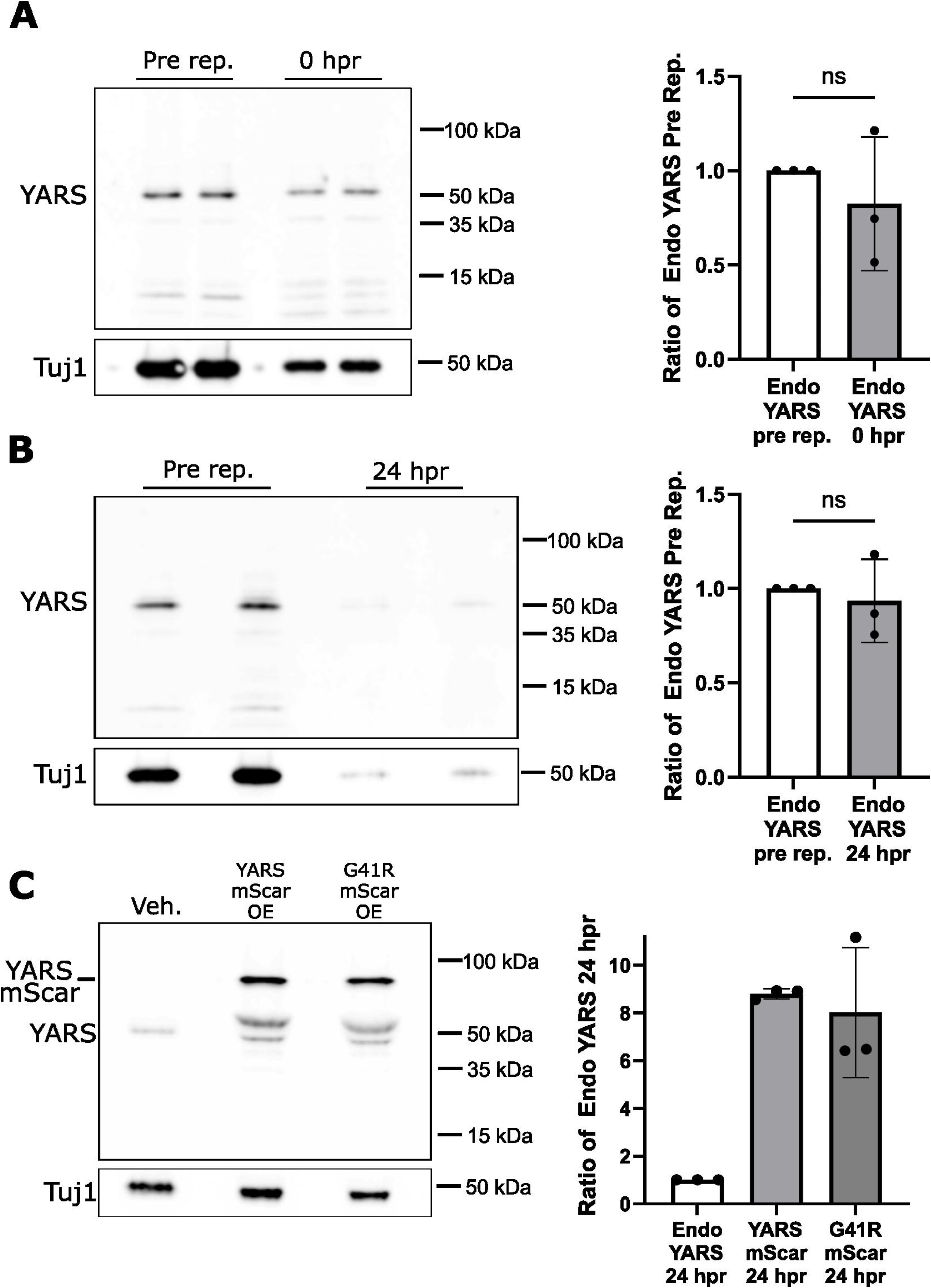
Replating does not alter expression of endogenous or exogenously expressed YARS proteins. **(A)** Whole cell extracts were collected from neurons on DIV8 as naïve controls or immediately after replating (0hr). Endogenous YARS protein levels were detected by western immunoblotting and normalized to Tuj1 as a load control. Two replicates are shown in the representative western blot from both conditions. **(B)** We performed a similar comparison twenty-four hours post-replating (24hpr). In this experiment we detected a noticeable decrease in endogenous YARS protein however a comparable decrease was measured in Tuj1 protein from these replicate samples. In **(C)** neurons were transduced with lentiviruses expressing YARS-mScarlet (WT or G41R) and protein levels evaluated from samples collected pre-twenty-four hours post-replating. Ratio to endogenous Yars was quantified and shown in graphs on the right. Error bars represent +/-1 SD. We performed a one sample Wilcoxon test in **(A) & (B)**. ns. not significant.

We also evaluated protein levels of exogenously expressed YARS. DRGs were transduced with a version of WT-YARS and G41R-YARS tagged with mScarlet to separate overexpressed YARS levels with endogenous YARS on a western blot. WT- YARS and G41R expression levels were nearly eight-fold higher than endogenous YARS in lysates collected 24hpr after replating when protein synthesis defects are substantially diminished (Figure 6*D*), similar to expression levels observed in non-replated DRGs (Supplementary Figure 1C). Therefore, replating does not rescue protein synthesis through increasing endogenous YARS or reducing exogenously expressed YARS.

### Replating-induced rescue of protein synthesis requires mTOR

Axotomy stimulates regeneration in PNS neurons and we predicted that severing distal axons would also rescue protein synthesis. We performed axotomy in non-replated DRG neurons expressing G41R-YARS at DIV 8; however, severing axons was not sufficient to rescue protein synthesis (Supplementary Fig. 3). Replating stimulates transcriptional upregulation of regeneration-associated genes (RAGs) (Mahar and Cavalli, 2018; Tome and Almeida, 2024). To test whether new transcription is required for rescuing protein synthesis in G41R neurons, we applied the transcription inhibitor actinomycin D to replated DRGs and measured TS fluorescence over the next 12 hours.

In the presence of actinomycin D, TS synthesis was delayed in G41R-expressing neurons at early timepoints after danoprevir addition, (Fig. 7*A*) however no statistically significant differences were noted at the twelve-hour time point (Fig. 7*B*). Actinomycin D treatment did not reduce TS fluorescence in empty vector or WT-YARS conditions suggesting reduced synthesis in G41R-YARS was not due to a global impairment in TS synthesis triggered by inhibiting transcription. These results indicate new transcription contributes to the restoration of protein synthesis in G41R-YARS neurons after replating, though other factors likely participate in this effect.

**Figure 7.**
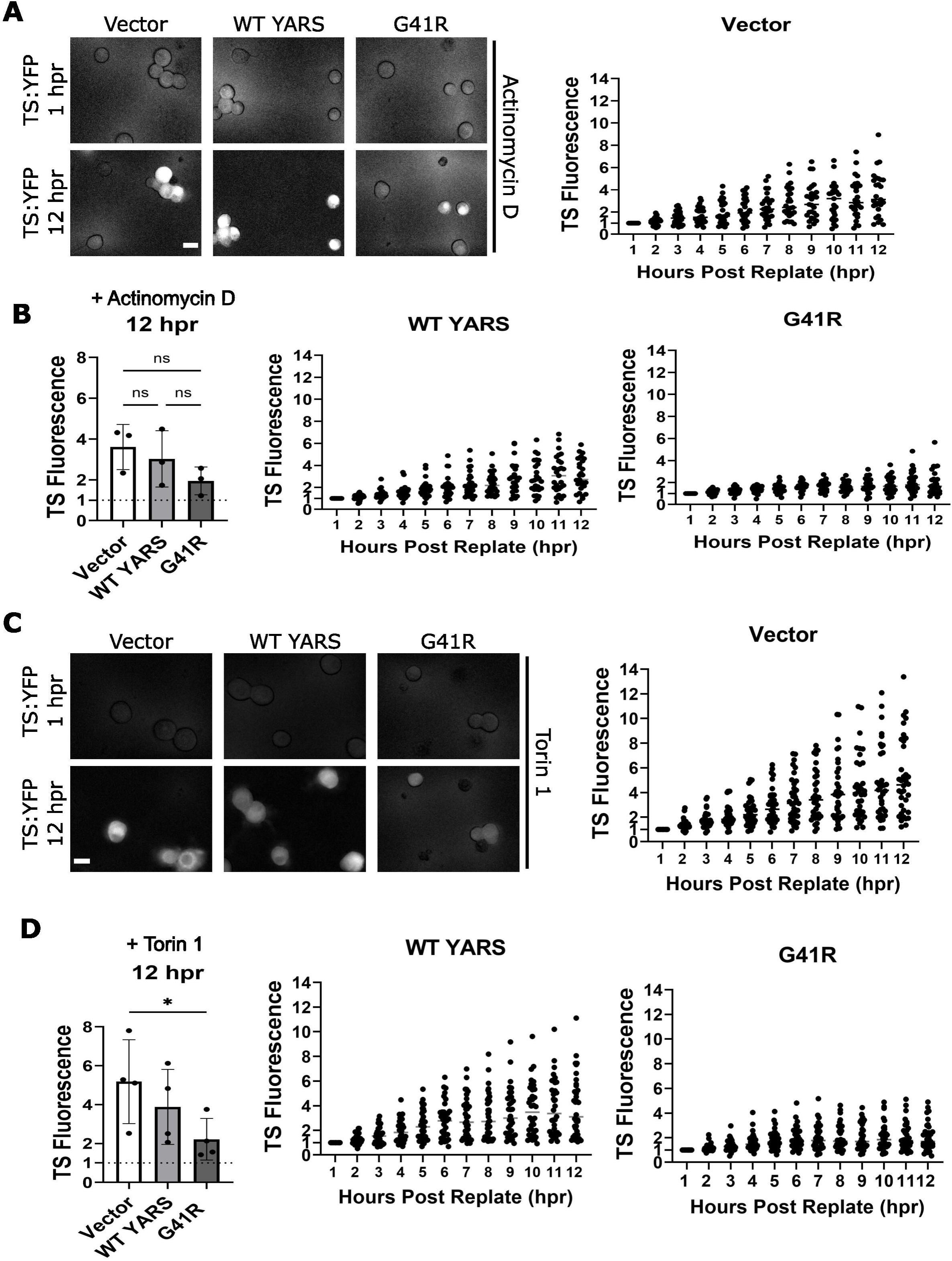
New transcription and mTOR participate in rescue of protein synthesis in replated neurons. **(A)** Replated neurons were pretreated with the transcription inhibitor actinomycin D (1µg/mL) then TS fluorescence measured after danoprevir application. **(B)** Change in TS fluorescence in individual cells through twelve hours post application (N= 3). **(C)** Replated neurons were pretreated with Torin 1 (150 nM) then TS fluorescence measured after danoprevir application. **(D)** Change in TS fluorescence in individual cells through twelve hours post application (N= 4). Error bars represent +/-1 SD. For statistical tests, one-way ANOVA was performed with post-hoc t-tests where *p<0.05. ns. not significant.

Axon injury stimulates mammalian target of rapamycin (mTOR), a well-established regulator of protein synthesis that promotes axon regeneration (Park et al., 2008; Abe et al., 2010; Terenzio et al., 2018; Koley et al., 2019). We treated neurons with Torin 1, an inhibitor of both mTORC1 and mTORC2 after replating. Torin1 treatment caused a reduction in TS synthesis in replated G41R-expressing neurons compared to WT or empty vector conditions (Fig. 7*C* and *D*). Therefore, the mTOR pathway is necessary for rescuing protein synthesis in YARS-G41R neurons after replating.

## DISCUSSION

AaRS supply amino acid-charged tRNAs for new protein synthesis and perturbing any one of these enzymes can have devastating consequences. For example, homozygous recessive mutations in YARS cause multisystem failure (Nowaczyk et al., 2017) while autosomal dominant mutations manifest in DI-CMTC, a peripheral neuropathy restricted to sensory and motor neurons (Jordanova et al., 2003; Jordanova et al., 2006; Hyun et al., 2014; Gonzaga-Jauregui et al., 2015). The mechanistic basis for such specificity has been a recurring question. Moreover, since CMT symptoms often emerge in early adolescence or later, how do PNS neurons successfully innervate their targets during development when the need for new protein synthesis would be especially high? We propose that upregulation of axon outgrowth pathways during PNS development insulates these neurons against CMT-YARS and the switch from development to maintenance diminishes these safeguards.

Our sensory neuron model recapitulates several key observations noted in other CMT models including reduced protein synthesis and a role for the ISR in neurodegeneration. Caspase-dependent cell death is not reported in mouse or drosophila models of YARS-CMT. Prolonged repression of protein synthesis might be more severe in our model to the extent of triggering apoptotic cell death. We bypassed this limitation through Bcl-xL overexpression which prevents cytochrome c release and caspase activation. Under these conditions, sensory neurons tolerate prolonged CMT- YARS expression over eight days in culture enabling us to evaluate long-term impacts on protein synthesis and conduct replating studies for axon regrowth.

Neurons expressing three different YARS alleles identified in patients with DI- CMTC reduced protein synthesis, confirming prior studies with two alleles (G41R and del153) and testing D81I for the first time. Poor expression precluded us from evaluating E196K which shows minimal loss of enzymatic activity *in vitro* yet does elicit reduced protein synthesis and neuropathy *in vivo* (Froelich and First, 2011; Niehues et al., 2015). Protein synthesis defects preceded caspase activation and were not suppressed by Bcl- xL overexpression, suggesting CMT-YARS expression triggers a decline in protein synthesis upstream of caspase activation. Our findings do not rule out gain-of-function interactions as the mechanism underlying CMT-YARS neuropathy (Blocquel et al., 2017; Bervoets et al., 2019; Ermanoska et al., 2023) which might repress protein synthesis independent of aminoacylation activity. Regardless, diminished protein synthesis would hinder neuronal function and compromise health due to depletion of short-lived proteins.

PNS cells maintain the capacity for axon regeneration into adulthood. Nerve transection activates local mTOR signaling which boosts synthesis of pro-regenerative factors (Tome and Almeida, 2024). Retrograde signals from the lesion site stimulate transcription of RAGs which collectively drive new axon outgrowth. mTOR activation is the most direct route toward stimulating protein synthesis however axotomy was not sufficient to rescue protein synthesis in YARS-G41R neurons. Additional signaling events activated during the replating procedure are likely required for synthesis rescue. Trypsin facilitates substrate detachment and might stimulate intracellular signaling through cleavage of a transmembrane receptor. Axon regrowth after replating is highly branched compared to regrowth after axotomy which indicates loss of polarity or activation of a distinct gene repertoire. The role of new transcription supports the latter and might involve upregulation of tRNAs which sequester CMT-aaRS and reduce neuropathy in animal models (Zuko et al., 2021). Identifying the minimal stimulus necessary to restore protein synthesis will open new avenues for treating this neuropathy.

Developing neurons are remarkably resistant to pathological stressors that manifest in adulthood, suggesting a decline axon outgrowth inversely correlates with resilience against proteotoxic threats. Protein synthesis tapered off in YARS-G41R neurons four days after replating, consistent with the gradual reduction of regenerative markers in replated neurons (Frey et al., 2015) and slower axon growth. Boosting regenerative capacity to repair the nervous system has been a long-term aspiration in neuroscience. Our study reinforces the value of this ambitious goal as a therapeutic option for CMT and other neurological disorders.

## METHODS

### Plasmids and Reagents

TimeStamp (TS) was amplified from PSD95-TS-YFP (Addgene Plasmid #43335; http://n2taddgene:4225; RRID:Addgene_42225) and subcloned downstream of the human ubiquitin promoter with Gibson Cloning. Human tyrosyl tRNA synthase (YARS) and variants were subcloned downstream of the human ubiquitin promoter with Gibson cloning. Sequences for generating sgRNAs targeting mouse Bax were #1 5’ GTTTCATCCAGGATCGAGCA3’ and #2 5’ TTGCTGATGGCAACTTCAAC 3’. Scramble sgRNA sequences, Cas9 expression plasmid, Bcl-xL, and myristoylated mScarlet were used as previously described (Danos et al., 2025). Antibodies for western immunoblotting were used as follows: anti-Yars (Bethyl Laboratories; RRID: AB_2631459; 1:1000), anti-Tuj1 (BioLegend; RRID: AB_2562570; 1:10,000), anti-Gapdh (Santa Cruz Biotechnology; RRID: AB_10847862; 1:500), and anti-GFP (Thermo Fisher Scientific; RRID: AB_221569). Chemicals were from the following vendors. Torin 1, danoprevir, ISRIB trans-isomer, and GCN2IB were from Medchemexpress. Actinomycin D, 8-CPT-Cyclic AMP, and anisomycin were from Cayman Chemical. Chemicals were prepared as a stock solution and stored as aliquots per recommendations from the vendor. Each aliquot was used once and discarded.

### Culture of primary sensory neurons

All mouse procedures were reviewed and approved by the University of Iowa Office of Institutional Animal Care and Use Committee. Timed pregnant mice were purchased from Charles River Laboratory. Dorsal root ganglia (DRG) were dissected from E13.5 mouse embryos and seeded on plates coated with poly-d-lysine and laminin. DRG sensory neurons were incubated in Neurobasal media (Gibco) supplemented with B27 (Gibco), 50ng/mL recombinant beta nerve growth factor (Proteintech), and 1mM 5- fluorodeoxyuridine/1mM uridine (Thermo Fisher Scientific) to eliminate mitotic, non-neuronal cells. Neurons were maintained at 37°C and 5% CO^2^ for the duration of each experiment.

### Cell Lysis and western immunoblotting

DRGs were lysed in RIPA buffer (50mM Tris-HCl pH 7.4, 1mM EDTA, 1% Triton X-100, 0.5% sodium deoxycholate, 0.1% sodium dodecyl sulfate, and150mM NaCl) supplemented with phosphatase inhibitors and protease inhibitors (Thermo Scientific). Cell extracts were centrifuged at 5,000xg for 5min to pellet debris and the supernatant added to Laemmli buffer with fresh β-mercaptoenthanol. Samples were separated by denaturing polyacrylamide gel electrophoresis and transferred onto nitrocellulose for western immunoblotting with antibodies listed above.

### TimeStamp analysis of new protein synthesis

DRGs seeded in 96-well plates were transduced with the lentiviruses expressing the following on DIV2: TimeStamp (TS), Bcl-xL, myristoylated mScarlet, and YARS-CMT variants. On DIV7 cells were treated with 1µM danoprevir to inhibit NS3 protease activity in the TS reporter and fluorescence images collected with an automated microscope (either Cytation 5 or Lionheart) every hour for thirteen hours. Analysis was performed with ImageJ. Briefly, timelapse images from the same field were assembled into a stack. We performed rolling ball background subtraction then fluorescence intensity quantified from individual cells for each timepoint. TS fluorescence for individual cells was calculated at each timepoint as a ratio of fluorescence intensity measured from that cell at 1 hour post danoprevir addition because of a shift in the visual plane during image acquisition from 0hr to 1hr. We quantified TS fluorescence from at least 10 cells per experimental replicate (from different wells) and at least 30 total cells from DRG preparations generated from independent mouse litters.

### Detection of activated caspase3/7

Caspase activity was visualized with CellEvent™ Caspase-3/7 detection reagent (Invitrogen) following manufacturer’s suggestions. Briefly, DRG neurons expressing G41R YARS or a control were incubated with the Caspase 3/7 detection reagent on DIV 4 or DIV 6 then images collected with an automated microscope over a twelve-hour period. We noticed that caspase positivity would be transient and often disappear when the plasma membrane ruptured which could lead to false negative findings. To circumvent this technical limitation, we counted the total number of positive cells observed this twelve-hour interval and report this as % caspase positive from an experimental condition.

### O-propargyl-puromycin (OPP) labeling of newly synthesized protein

DRG neurons seeded in 96-well dishes were transduced with lentivirus expressing myristoylated mScarlet and Bcl-xL on DIV 2. Lentivirus expressing the CMT-YARS or an empty vector were applied on DIV 4. We labeled newly synthesized proteins using reagents and procedures from an OPP Protein Synthesis Assay kit from Vector Laboratories with the following modifications. On DIV 8 we applied OPP by performing a half-media change with fresh media containing OPP (20µM). Some wells underwent a media change with fresh media lacking OPP to determine background. Other positive control wells were pretreated with 25µg/mL cycloheximide. Cells were incubated with OPP-containing media for ten minutes, then media was replaced once with cold PBS and again with cold PBS plus 3.7% formaldehyde. Click chemistry was performed following manufacturer’s recommendations and images collected with a Lionheart FX. Fluorescence intensity was measured from at least fifty cells from three wells per experimental replicate.

### Replating DRGs for analysis of axon outgrowth and TS fluorescence

DRGs were densely seeded on 12-well plates. On DIV 2, DRGs were transduced with lentiviruses expressing myristoylated mScarlet, Bcl-xL and conditional YARS lentivirus (empty vector, WT-YARS, G41R-YARS). For proteins synthesis studies TS lentivirus was added on DIV 5. On DIV 8, cells were dissociated from the plate with trypsin (0.05% EDTA-Gibco), triturated with a plastic pipette, washed in normal media, and seeded onto a new plate. For regrowth and TimeStamp studies, cells were seeded in 96-well plates. For protein biochemistry, cells were immediately lysed or seeded in 12- well plates and lysed twenty-four hours later. For regrowth studies, myristoylated mScarlet images were collected every hour in an automated microscope. Images were analyzed with a custom ImageJ macro that auto-contrasts and thresholds each image then measures axon particles as total axon area (TAA) from the binarized image. This procedure eliminates most signal from the cell body due to high focal fluorescence.

Axon regrowth was visualized over eleven hours with automated microscopy and quantified by measuring total axon area at a given timepoint normalized to total axon area at time 1hr. At least ten image fields measured over time per condition per replicate. For each timepoint, the TAA in a given image field was calculated as a ratio of time 1hr TAA from that field. Axonal regrowth measurements for at least thirty fields per condition were calculated this way and averaged respective to experimental replicate. In pharmacology studies, drugs were applied immediately after replating and left in the media during the duration of the experiment. Protein synthesis experiments were conducted with the TS reporter following procedures described above.

## Data analysis and statistics

Graphs and statistical analysis were performed in GraphPad Prism. Specific statistical tests employed for each experiment are described with the accompanying figure legend. Western immunoblots were quantified in ImageJ with signals normalized internally to the Tuj1 load control. Experimental replicates were generated from sensory neuron cultures derived from independent mouse litters and at least two independent lentiviral preparations.

## Acknowledgements and Funding

Research conducted in this manuscript was supported by funds from the National Institutes of Health to D.W.S (R01NS127781).

## Author Contributions

J.K.; conceptualization, data generation, data analysis, and writing. A.M., Y.H. and J.A.; data generation. D.W.S. conceptualization, data generation, data analysis, and writing.

---

**Supplementary Figure 1.**
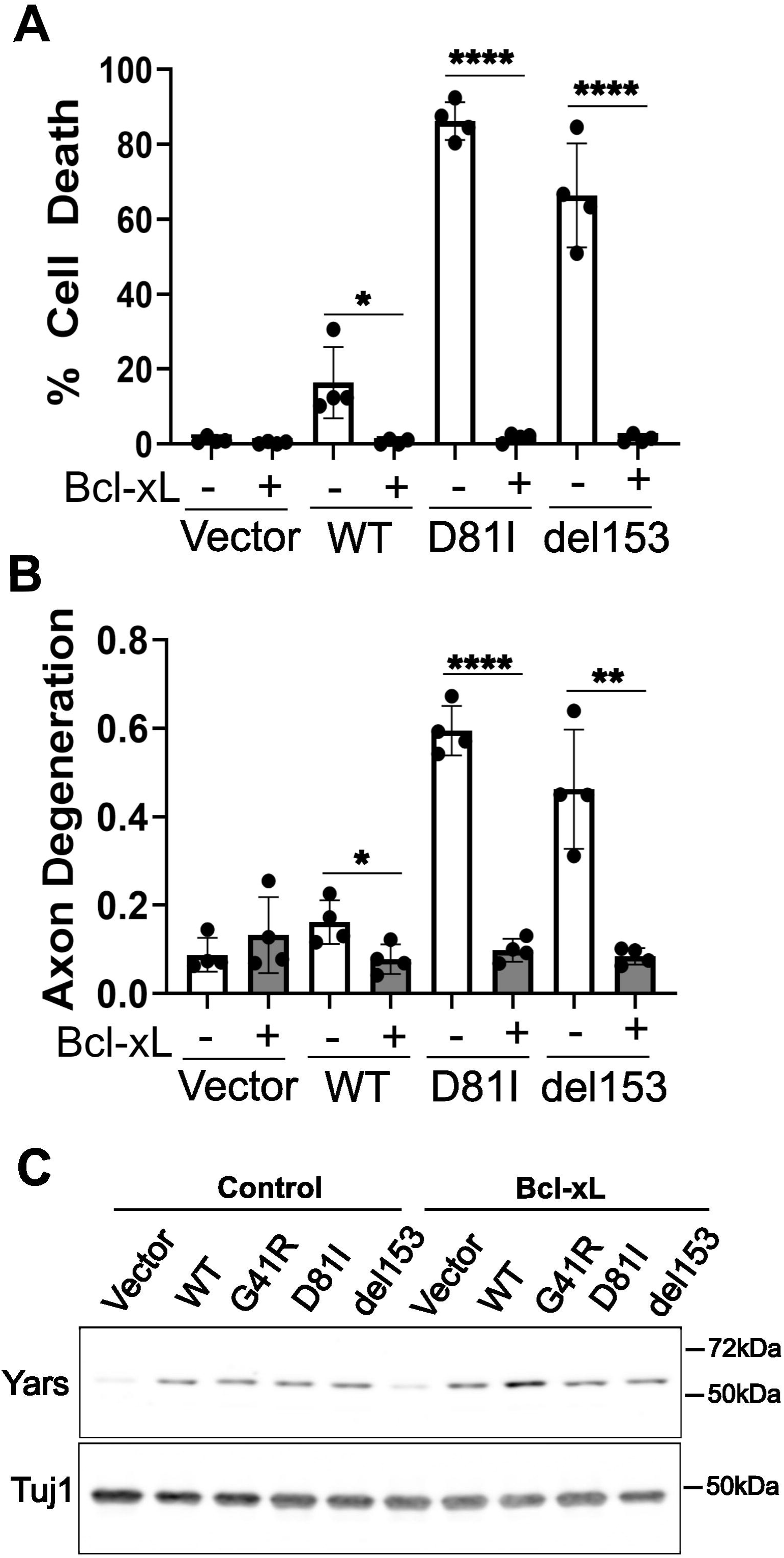
Caspase-dependent axon degeneration and cell death in DRG neurons expressing YARS D81I or YARS del153. **(A)** Cell death and **(B)** axon degeneration quantified from DRG neurons expressing an empty vector, wildtype YARS or, the indicated CMT-YARS protein in the presence or absence of Bcl-xL overexpression (N=4). **(C)** Bcl-xL expression does not affect CMT-YARS protein levels in DRGs. Error bars represent +/-1 SD. For statistical tests, one-way ANOVA was performed with post-hoc t-tests where *p<0.05, ** p<0.01, and ****p<0.001.

**Supplementary Figure 2.**
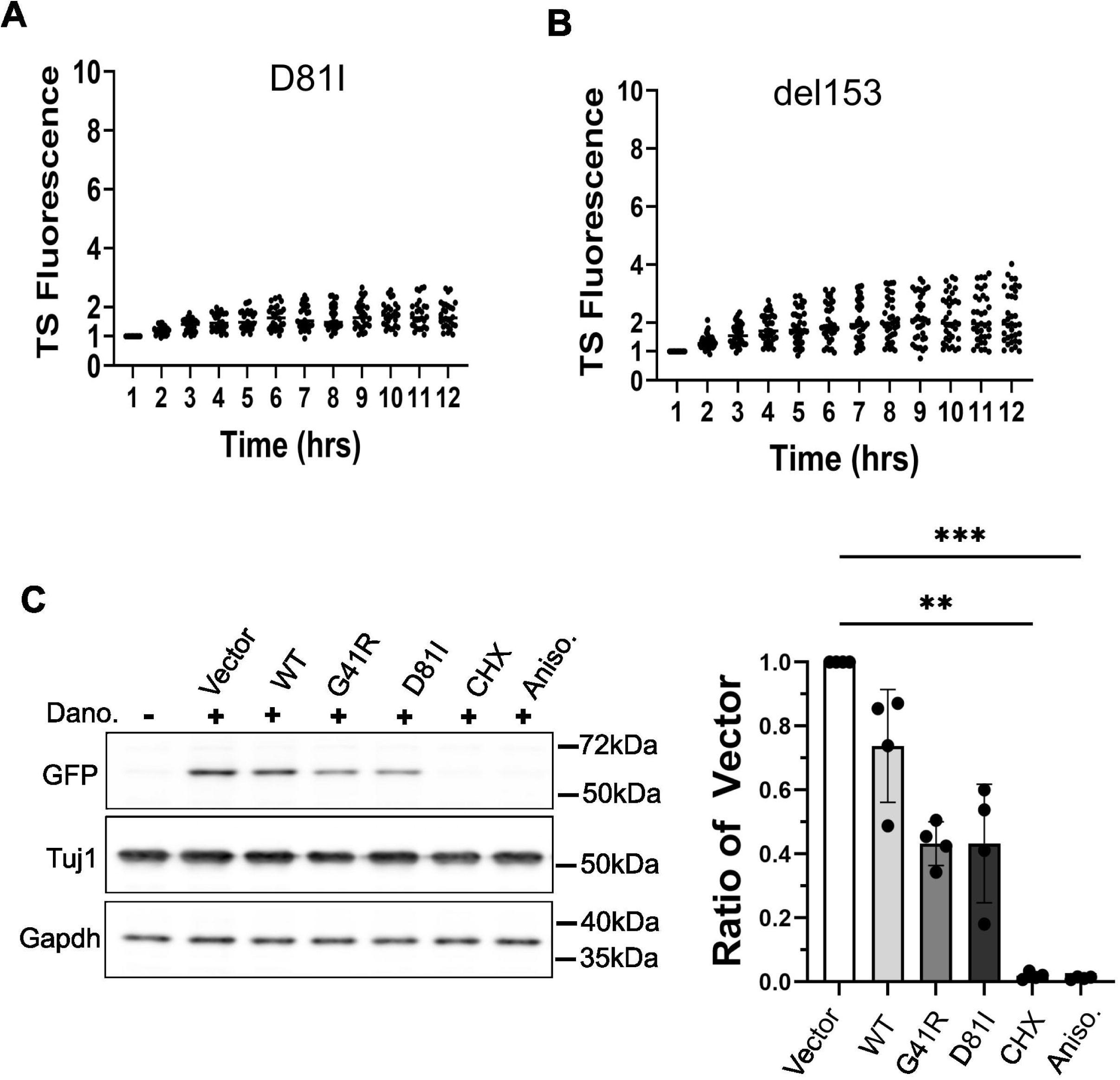
Reduced protein synthesis in sensory neurons expressing CMT-YARS. **(A) & (B)** TS fluorescence from individual cells after danoprevir addition expressing D81I-YARS (28 cells from three independent experiments) or del153-YARS (36 cells from 3 independent experiments). **(C)** Western blot of DRG extracts four hours after danoprevir addition. TS was detected with an antibody to GFP = with quantification on the right (N=4). Error bars represent +/-1 SD. We performed a Kruskal-Wallis analysis with Dunn’s post-hoc test to assess significance where ** p<0.01, and ***p<0.005.

**Supplementary Figure 3.**
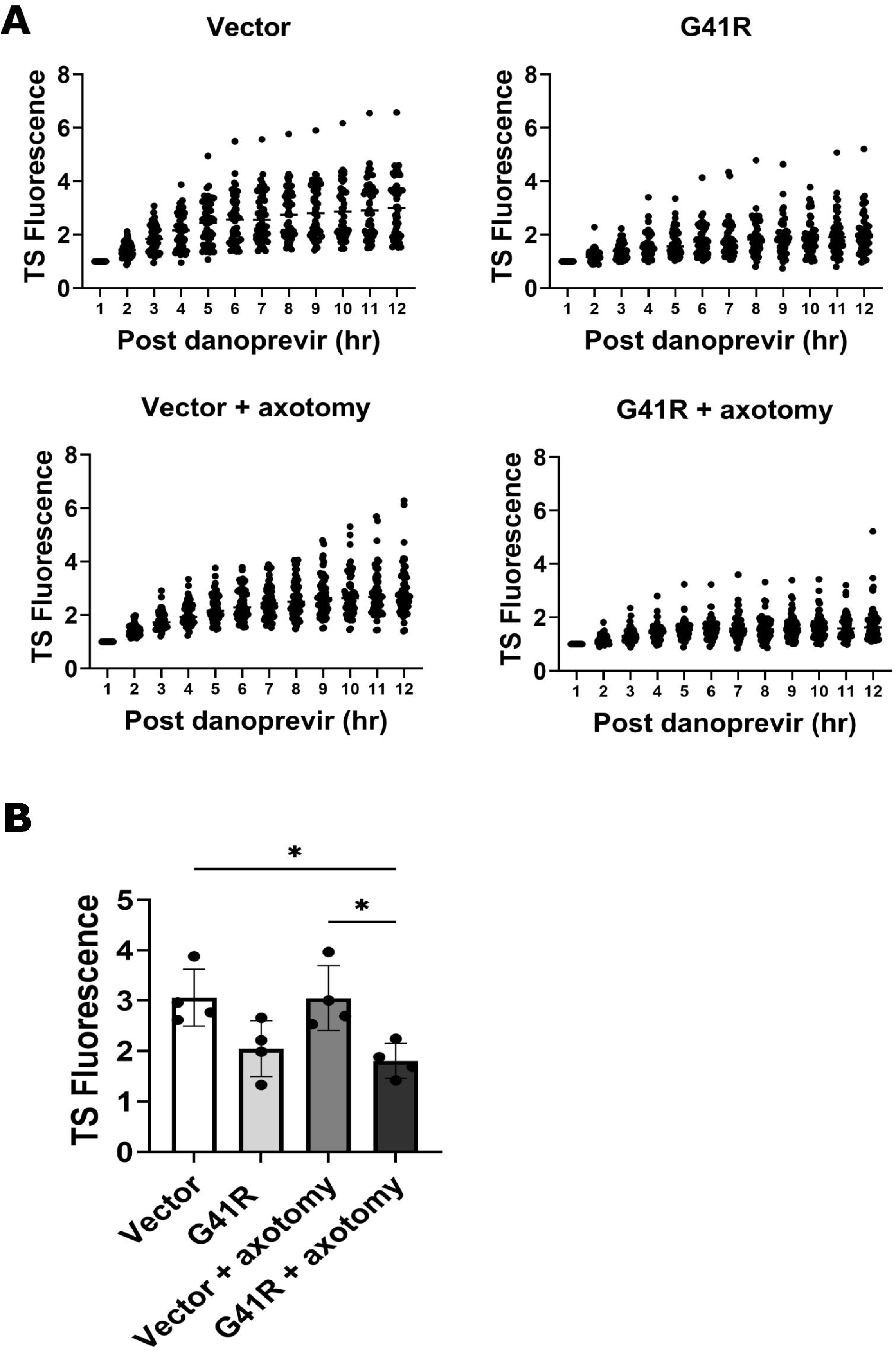
Axotomy is not sufficient to restore protein synthesis in YARS-G41R sensory neurons. **(A)** DRG sensory neurons seeded in a spot culture were transduced on DIV2 with lentivirus expressing an empty vector or YARS-G41R, Bcl-xL to prevent caspase activation, and TimeSTAMP. On DIV8 a razor was used to sever axons around the spot culture. One hour later, danoprevir was added and TimeSTAMP visualized over the next twelve hours. Data points from individual cells are shown for each condition (from at least 40 cells per condition in four independent replicates) with the twelve-hour time point **(B)**. Error bars represent +/-1 SD. For statistical tests, one-way ANOVA was performed with post-hoc t-tests where *p<0.05.

